# Disrupted dynamic network reconfiguration of the executive and reward networks in internet gaming disorder: Why IGD subjects cannot control their gaming cravings

**DOI:** 10.1101/2021.09.29.462308

**Authors:** Min Wang, Hui Zheng, Weiran Zhou, Bo Yang, Lingxiao Wang, Shuaiyu Chen, Guang-Heng Dong

## Abstract

**Background:** Studies have shown that people with internet gaming disorder (IGD) show impaired executive control over their gaming cravings; however, the neural mechanisms underlying this process remain unknown. In addition, these conclusions were based on the hypothesis that brain networks were temporally stationary and neglected changes in cognitive processes.

**Methods:** Resting-state fMRI data were collected from 402 subjects (162 subjects with IGD and 240 recreational game users (RGUs)). The community structure (recruitment and integration) of the executive control network and the basal ganglia network (BGN, representing the reward network) of patients with IGD and healthy controls were analyzed and compared. Mediation effects were analyzed among the different networks.

**Results:** When compared to RGUs, subjects with IGD had a lower recruitment coefficient within the right executive control network (ECN). Further analysis showed that only male subjects had a lower recruitment coefficient. Mediation analysis showed that the integration coefficient of the right ECN mediated the relationship between the recruitment coefficients of both the right ECN and the BGN in RGUs.

**Conclusions:** Subjects with IGD had a lower recruitment coefficient than RGUs, and this feature was only observed in male subjects, making them less efficient at impulse control. The meditation results suggest a top-down control mechanism of the ECN is missing in subjects with IGD. All of these findings could explain why subjects with IGD have impaired executive control over their gaming cravings.

## Introduction

Many studies have proven that people with internet gaming disorder (IGD) have impaired executive control over their gaming cravings ^1-3^. First, studies using different paradigms have shown impaired brain activation in executive control-related brain regions ^4^, including the dorsolateral prefrontal cortex (DLPFC) and the orbitofrontal cortex ^5,6^. At the same time, subjects with IGD had increased gaming cravings, which could be detected by increased brain activity in reward processing-related brain regions ^7-9^. Second, abnormal functional connectivity (FC) between brain regions involved in executive control and reward processing has been reported in subjects with IGD ^10,11^. People with IGD had lower FC between the orbitofrontal cortex and the dorsal striatum ^12,13^, the left medial orbitofrontal cortex and the putamen, and the DLPFC and the putamen ^14,15^. A study with a large sample size (337 subjects) observed that subjects with IGD had lower FC between the middle frontal gyrus and the putamen, inferior frontal gyrus and ventral striatum. Disorder severity and craving scores were negatively correlated with FC between striatal and frontal brain regions ^16^.

According to an increasing number of studies, cognitive function is accomplished by the interactions of a brain network rather than by specific regions ^17,18^. Current studies suggest that there are two distinct brain networks that jointly influence our choices: the executive control network, which is associated with inhibiting impulses, and the reward network, which mediates immediate rewards ^19,20^. Abnormal interactions between these two networks were observed in drug-addicted groups ^20^, heroin-dependent subjects ^21^, and subject with IGD^10^, which suggests that impairments in executive control led to ineffective inhibition of enhanced cravings for excessive online game playing. This can shed light on the mechanistic understanding of addictive behaviors at the large-scale system level. The combination of enhanced seeking motivations and an inability to inhibit impulsive behaviors is thought to represent a failure of executive control ^22,23^.

Limitations existed in these studies. First, all studies described the current features of patients with IGD having impaired executive control over cravings; however, none of them could explain why subjects with IGD have disturbed processes during this process, and the neural mechanisms underlying this feature remain unclear ^2,24,25^. Second, these researchers hypothesized that the interactive connection between brain regions/networks throughout the process was temporally stationary ^26-28^. However, the human brain is a complex system; the active states and functional connectivity of the human brain can change during learning, growth, and even rest ^29,30^. The brain dynamically integrates and coordinates the interaction of different brain areas to complete complex cognitive functions.

Thus, to fully understand the neural foundations of impaired executive control over gaming cravings in subjects with IGD, it is necessary to explore the dynamic characteristics of the time-dependent changes in the executive control and reward networks. In fMRI scans, the blood oxygen level dependence could signal fluctuations in brain activity; thus, there is sufficient information to study the dynamic properties of brain networks ^31^. Several researchers have studied the variability of brain networks to detect dynamic functional changes, and they have demonstrated that it is a useful method for detecting dynamic changes in brain networks ^32^.

Community structure is a functionally relevant graph metric to study the organization and interaction of functional systems in the brain network ^33^. The interactive couplings within community nodes (or brain regions) are strong and dense, whereas interactive couplings between communities are sparse ^34^. The community structure (or modular organization) can delineate the functional segregation and integration of whole-brain networks. In most cases, two indices were used to measure the community features: recruitment, which refers to the probability that a brain region is in the same community as other nodes from its own network; and integration, which refers to the probability that a brain region is in the same community as nodes from other networks ^35^. Researchers have identified community structure in both structural and functional networks in the healthy human brain ^36^. Community structure has been used to identify potential functional modules for groups of brain regions exhibiting similar trajectories and functions over time ^36,37^. The way networks change over time can provide insights into neural mechanisms.

Recently, using community structure has piqued the interest of clinical researchers, as studies have proven that disrupted community structures are found in a variety of brain disorders ^38,39^. A study found a less stable community structure at the resting-state network level in a group of patients with schizophrenia and provided novel methods for exploring dynamic community structure ^40^. Lord et al. detected the community structure of the functional network for individuals with unipolar depression ^41^. Zheng et al. showed that, in major depression disorder (MDD), the anterior cingulate cortex exhibited abnormal flexibility in community structures ^42^ and that unmedicated MDD groups and medicated MDD groups exhibited a similar reconfiguration of the community structure of the visual association and the default mode systems, but that the groups had different reconfiguration profiles in the frontoparietal control subsystems ^43^. One study suggested that dynamic network measures may be an effective biomarker for detecting language dysfunction associated with neurological diseases like temporal lobe epilepsy ^35^.

The primary aim of the current study was to address the limitations of previous studies on IGD and provide a better understanding of the neural mechanisms underlying impaired executive control over cravings. To achieve this goal, we examined the distribution of community assignment across the entire scanning time and compared the changes in the community structure of the executive control network and reward network in subjects with IGD. We hypothesized that patients with IGD would have a different community structure than the healthy control group. Second, we wanted to explore how these community structure features affect the top-down controlling process and whether this feature can be observed in subjects with IGD. We hypothesized that the top-down control process is disrupted in subjects with IGD.

## Methods and procedures

### Participants

This study was approved by the ethics investigation committee of Hangzhou Normal University and conducted from Dec 2013 to Dec 2019. All participants signed written informed consent forms in accordance with the Declaration of Helsinki.

Four hundred thirty (430) online game players were recruited from universities in China. Twelve players with missing fMRI data and sixteen players with head motions > 2.5° during the scan were excluded. The remaining subjects were evaluated for the severity of IGD via both Young’s Internet Addiction Test (IAT) (www.netaddiction.net) and DSM-5 diagnostic criteria ^44^. Individuals with IGD had to have IAT scores greater than 50, as well as at least five of the nine DSM-5 criteria (as they were recruited before the release of DSM-5, 41 subjects from Dec 2013 did not have DSM-5 data). A total of 162 subjects were diagnosed with IGD, and the remaining 240 were classified as recreational game users (RGUs) according to the assessment by trained researchers. For relevant demographic information, see Table 1. All subjects had no comorbid DSM-IV axis I or axis II disorders, based on the results of a structured psychiatric interview ^45^.

### Imaging acquisition

Resting-state functional data were acquired from a 3T Siemens Trio MRI system using an echo-planar imaging (EPI) pulse sequence(33 axial slices with 3 mm slice thickness, TR = 2000 ms, TE = 30 ms, matrix = 64 × 64, FOV = 220 × 220 mm, flip angle = 90°, and a total of 210 volumes). The structural images were from a high-resolution, T1-weighted magnetization-prepared rapid gradient echo sequence (TR = 1900 ms, TE = 2.52 ms, TI = 900 ms,, flip angle = 9°, matrix = 256 × 256, slices = 176, voxel size = 1 × 1 × 1 mm^3^).

### Image preprocessing

The DPABI version 5.2 toolbox was used to perform fMRI preprocessing ^46^. The first 10 volumes were removed for all data. Slice-time and motion parameters were then evaluated and corrected for the remaining volumes. For the data in individual space, spatial normalization with the EPI templates was carried out and transformed to the MNI space with a 2 × 2 × 2 mm^3^ voxel resolution. Normalized data were smoothed with a 6 FWHM Gaussian kernel. Finally, the obtained images were detrended and regressed as covariates with noisy signals from white matter (WM), cerebrospinal fluid (CSF) and head motion parameters (Friston 24 model).

### Network ROI selection

IGD is often accompanied by dysfunctions in frontostriatal circuits, which can be interpreted as a disconnection between the executive control network and the basal ganglia network. We used the Shirer et al. ^47^ atlas, which covers 14 resting-state networks, to define these functional networks a priori, and the left and right executive control networks (including 6 brain regions per network) and the basal ganglia network (5 regions) were extracted for the current analysis. The right executive control network and the basal ganglia network overlap in the right caudate nucleus and the middle frontal gyrus due to the functional diversity of the regions. The time correlation r value between each voxel within the overlapping area and the two networks was calculated using all subject data, and the right caudate and middle frontal gyrus were parcellated by using a “winner-takes-all” approach with a bootstrapping strategy ^48^ (see Fig. 1).

**Fig.1.**
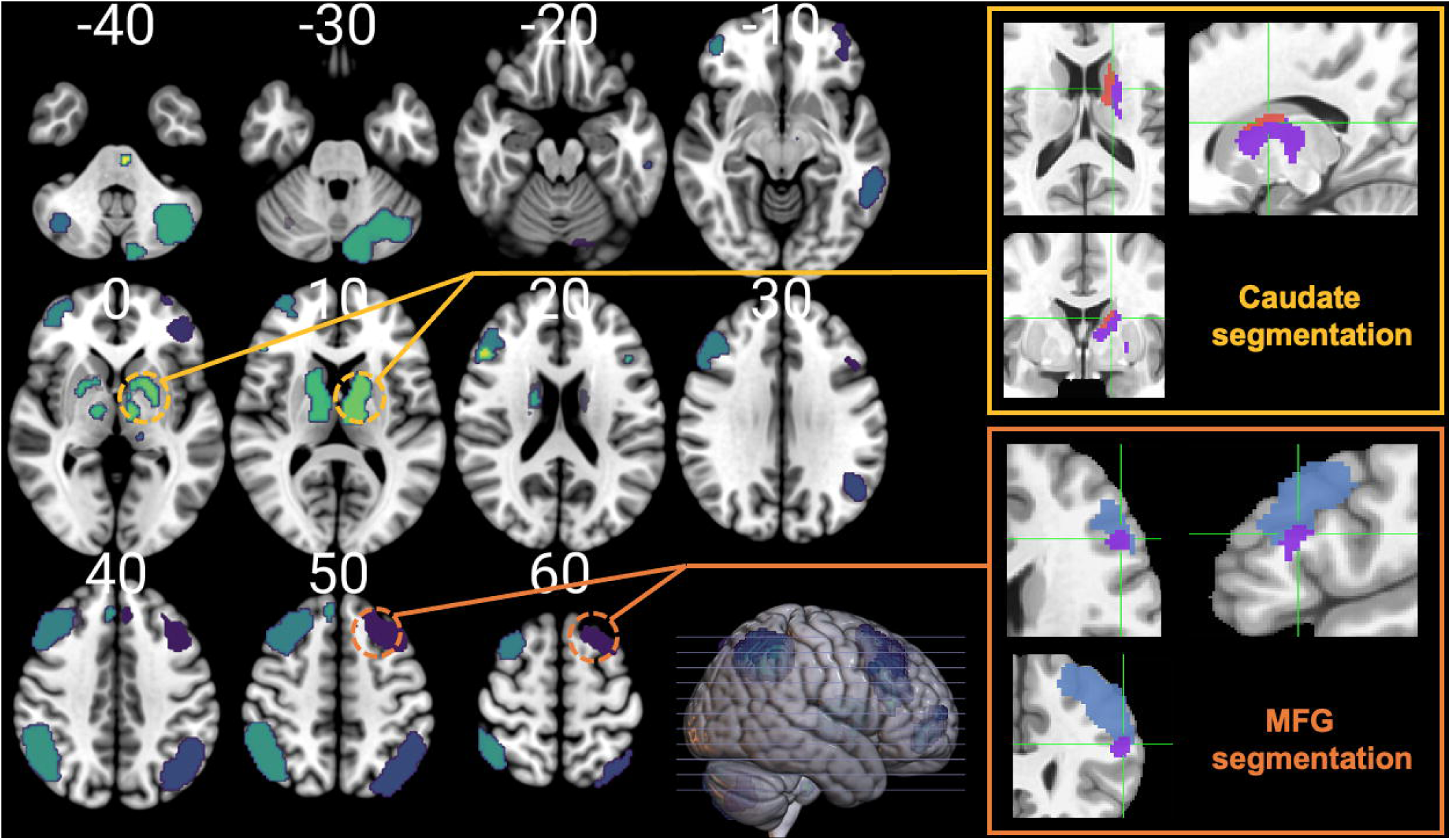
Segmentation of right caudate and middle frontal gyrus. These areas are divided into two networks based on the “winner-take-all” strategy. For the caudate, the area marked in pink is assigned as RECN, and the area marked in purple is assigned as BGN. For the middle frontal gyrus, the area marked in blue is assigned as RECN, and the area marked in purple is assigned as BGN.

### Network construction

We extracted the mean time series of 17 functionally defined regions of interest (ROIs). For network construction, we used wavelet decomposition rather than functional connectivity to extract information from these time series because it is more sensitive to subtle signal changes in nonstationary time series with noisy backgrounds ^49^. We use a maximal overlap discrete wavelet transformation with the Daubechies least asymmetric approach to decompose the time series into multiple frequency intervals ^50^: scale 1, 0.125∼0.25 Hz; scale 2, 0.062∼0.125 Hz; scale 3, 0.031∼0.062 Hz; and scale 4, 0.015∼0.031 Hz. Scale 3 was chosen as the main analysis frequency band because it is completely covered by the frequency range commonly interpreted in resting fMRI (i.e., 0.01∼0.1) and has higher sensitivity to disease classification ^51^.

The time-series data were then split into a consecutive series of 40 s time windows that overlapped with contiguous windows by 50%. The magnitude-squared spectral coherence between each pair of ROIs was estimated according to ^52,53^, generating a 17 × 17 adjacency matrix for each time window. Finally, the adjacency matrices in all 19 time windows would be linked together to form a multilayer network.

### Multilayer community detection

A community describes a group of nodes that are more strongly connected to each other than to nodes outside of their community ^54^, whereas a multilayer community further characterizes their reconfiguration over time. In the current study, we used a generalized Louvain community detection algorithm ^33^ involving the following multilayer modularity quality function:

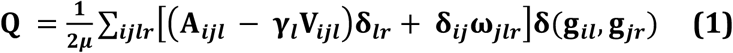

where *μ* is the total edge weight of the network, *A*_*ijl*_ is the edge between nodes *i* and *j* at layer *l* of the multilayer network, and *V*_*ijl*_ describes the corresponding element of a null model. The parameter *γ*_*l*_ sets the structural resolution parameter of layer *l* (i.e., the weight of intralayer edges) the parameter *ω*_*jlr*_ sets the temporal resolution parameter (i.e., the weight of interlayer edges, here *γ*_*l*_ = 1, *ω*_*jlr*_ = 0.4) ^55^, and the parameter *g* describes the community assignments of two nodes across the time domain, involving node *i* in layer *l* and node *j* in layer *r. δ* is a Kronecker delta function, where *δ(gil, gjr)* = 1 if *il* = *jr* and 0 otherwise.

Although the current network should be considered orderly and have interlayer links between sequential layers for nodes at the same position, the generalized Louvain algorithm has a stochastic nature, sometimes causing the instability of community assignments ^56^. To ensure the stability of the results, we performed 100 iterations for each subject and calculated the mean, similar to an implementation used in previous studies ^35^.

### Recruitment and integration

The 17 ROIs were categorized into three resting-state networks: the left and right executive control networks and the basal ganglia network (reward network). We calculated two dynamic indicators to quantify the dynamic interactions on inter- or intranetworks: recruitment and integration. The recruitment coefficient describes the average probability that node *i* is in the same community as other nodes from its own network and is defined as:

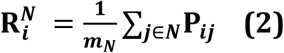

where *m*_*N*_ is the size of network *N*, calculated as the number of nodes in *N* and *P*_*ij*_ corresponds to the relative frequency at which nodes *i* and *j* were assigned to the same community across the time domain, where *Pij* = 1 if nodes *i* and *j* are always in the same community and 0 otherwise. Therefore, a node with high recruitment tends to be associated with nodes from its own network in the time domain. The integration coefficient describes the average probability that node *i* is in the same community as nodes from other networks, given by:

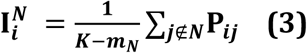

where *K* is the total number of nodes. A node with high integration tends to be associated with nodes from other networks in the time domain.

### Statistical analysis

Statistical analyses were conducted using SPSS 20.0. To match the two groups of subjects, we chose subjects from high to low according to their IAT scores. The first third of the subjects (134 subjects per group) were included. All statistical processes were performed at the network level, and significance was determined using independent sample t-tests with Bonferroni correction (*p <* .*016*). For the exploratory mediation analysis, PROCESS bootstrapping and bias-corrected 95% confidence intervals were used to assess the significance of the mediation model ^57^; a CI that does not contain 0 indicates a significant mediation effect. The analysis pipeline is depicted in Fig. 2.

**Fig.2.**
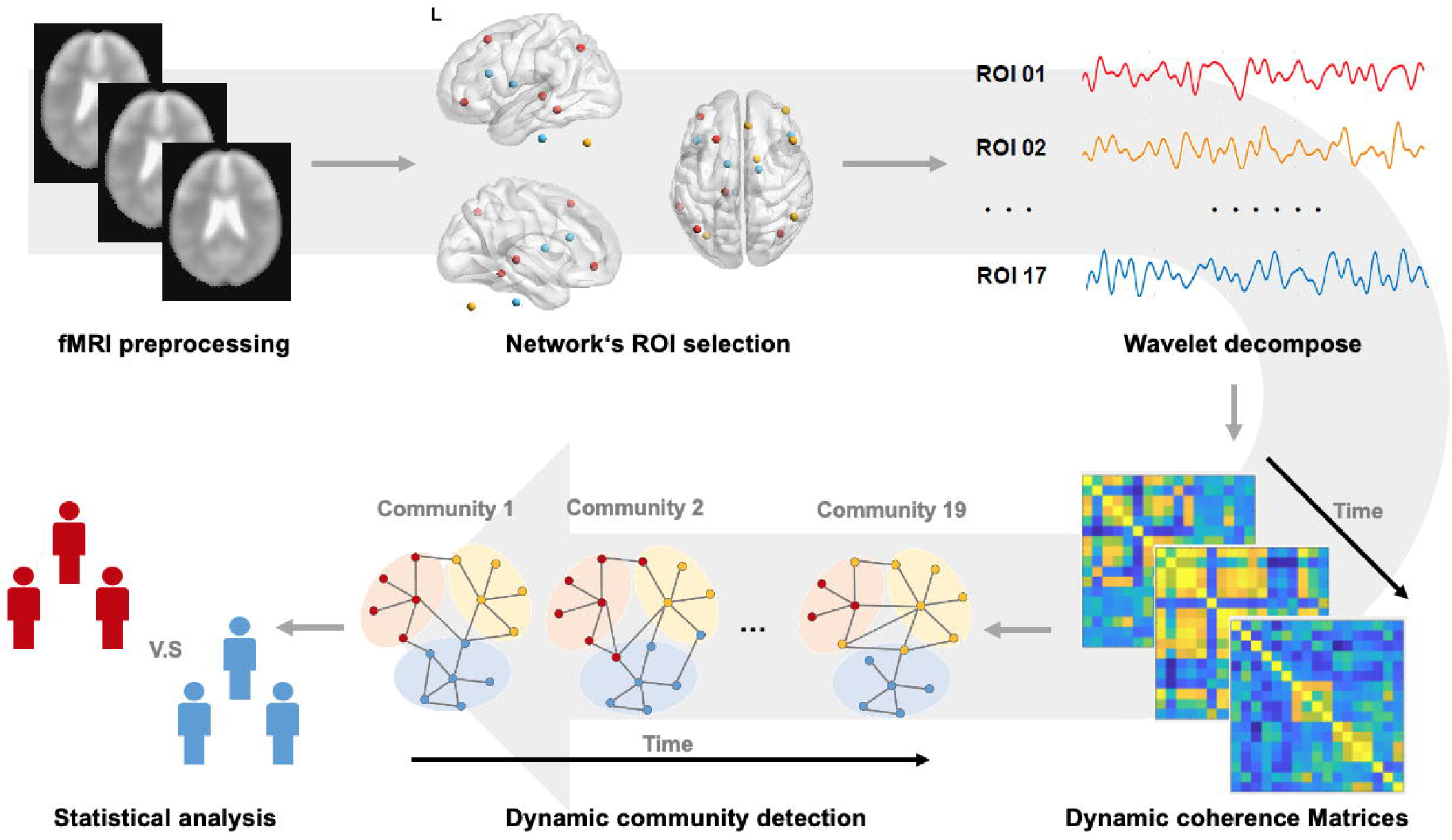
The analytic pipeline in current study. This figure shows the whole process from pre-processing to statistical analysis.

*Insert Fig. 2 here*

## Results

### Recruitment and integration

Compared to RGUs, subjects with IGD had a lower recruitment coefficient within RECNs (*t = -2*.*689, p = 0*.*007*, Bonferroni correction, see Fig. 3A). In general, a lower recruitment coefficient indicates that the nodes within the network are less likely to be divided into the same community over time. Thus, the current results might suggest that the functional network characterized by executive control in subjects with IGD was decoupled from the dynamic process. Although our hypothesis was associated with altered dynamic interactions between the ECN and BGN in subjects with IGD, no significant differences in the integration coefficient were found between the groups (Fig. 3B).

**Fig.3.**
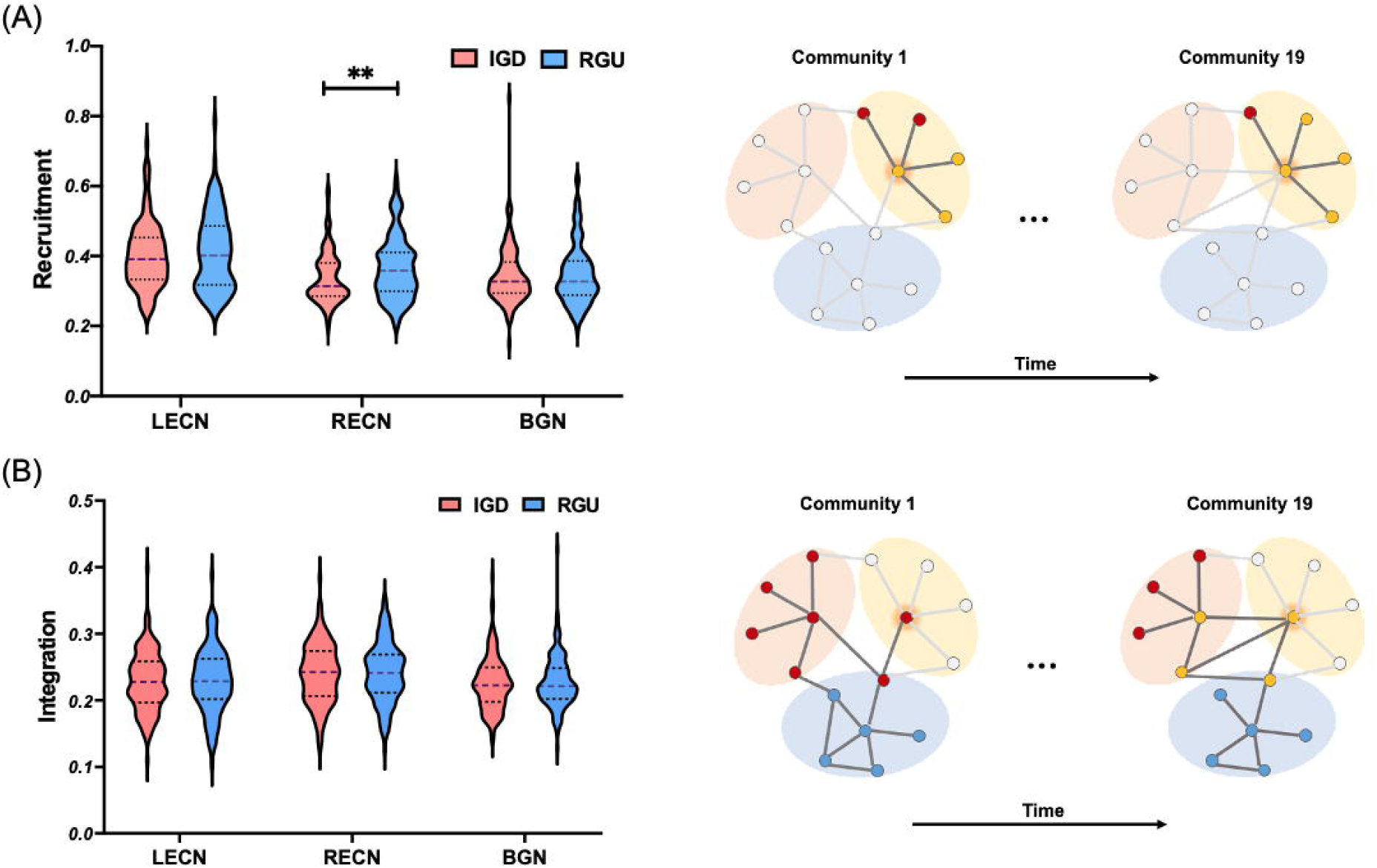
Between-group differences in recruitment and integration coefficients. Plot.A: left plot shows between-group difference in recruitment coefficient and right plot is a schematic diagram of the recruitment coefficient in a dynamic community; Plot.B: left plot shows between-group difference in integration coefficient and right plot is a schematic diagram of the integration coefficient in a dynamic community. **: *p* < 0.01.

In addition, we looked at the differences by sex in recruitment coefficients within RECNs. The results showed that there was a significant difference in the RECN recruitment coefficient only among males (*t = -2*.*467, p = 0*.*015)*, Bonferroni correction, see Fig. 4.

**Fig.4.**
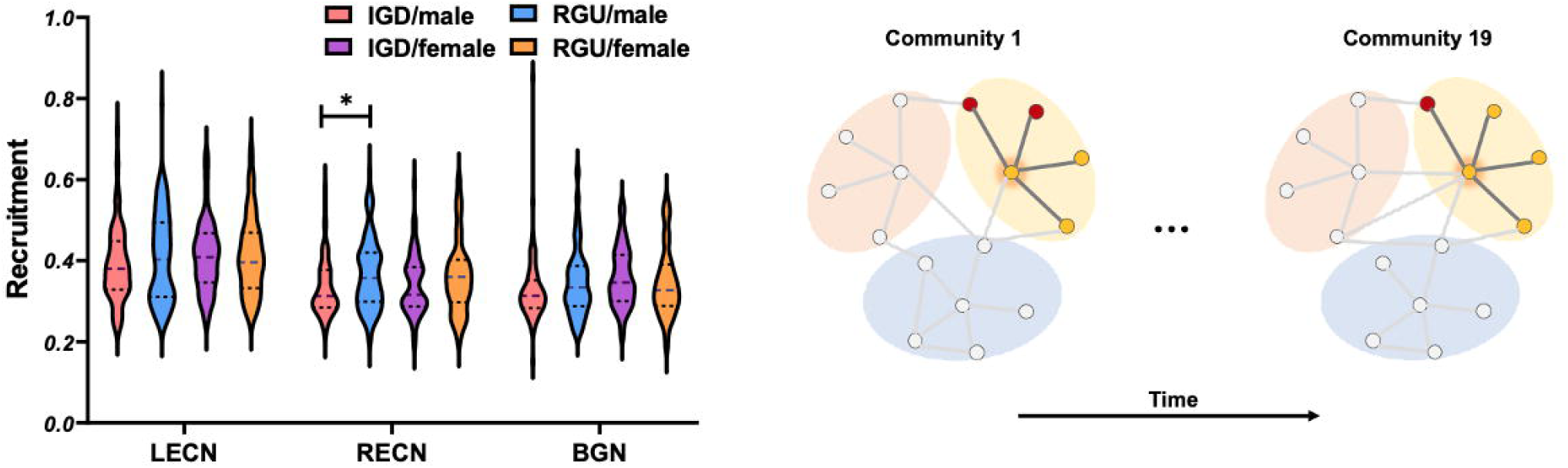
Between-group differences in recruitment coefficients based on sex. Left plot shows between-group difference in recruitment coefficient for males and females, while right plot is a schematic diagram of the recruitment coefficient in a dynamic community. *: *p* < 0.05.

### Mediation analysis

To further verify our hypothesis, an exploratory analysis of differences in the RCEN between groups was conducted by a mediation model. First, in the RGUs, we observed a two-way correlation of the integration coefficient of the RECN with both the recruitment coefficient of the RECN (*r = -0*.*225, p = 0*.*009*) and of the BGN (*r = 0*.*276, p = 0*.*001*), whereas no similar correlations were found in subjects with IGD. We then performed a mediation analysis using the integration coefficient of the RECN as the mediating factor, which showed that the integration coefficient of the RECN mediates the relationship between the recruitment coefficients of the RECN and BGN in RGUs (see Fig. 5B). The same model did not hold for subjects with IGD.

**Fig.5.**
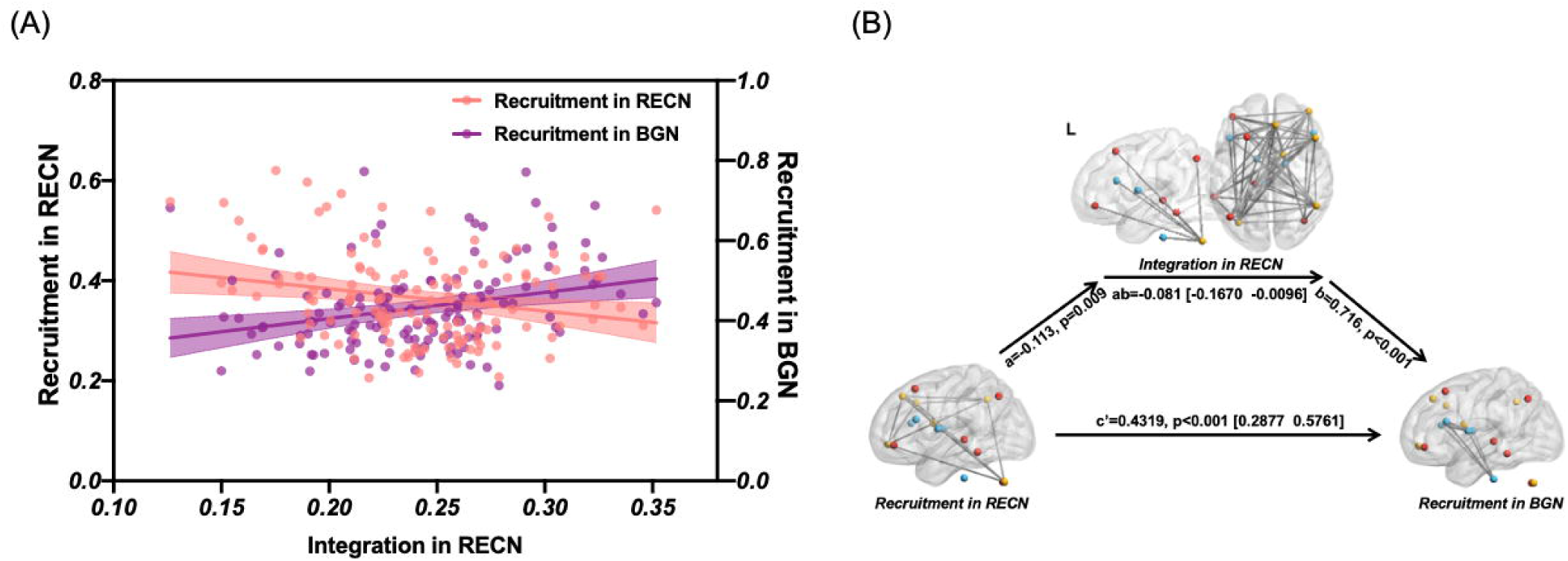
Mediation analysis. Left plot shows correlations of integration in RECN with the recruitment in BGN and RECN in RGUs, while right plot shows that integration in RECN significantly mediate the relationships between recruitment in BGN and RECN in RGUs.

### Statistical null models

We adopted the network null model to quantify the dynamic modular organization of the resting state network in subjects with IGD and RGUs ^52,58^. Descriptions and comparison results of these null models are provided in the Supplementary material. In short, the current results supported the dynamics of real network modules.

### Replication test with longitudinal data

To investigate further, we tracked 40 subjects (22 IGD, 18 RGU) (these 40 subjects were part of the 402 subjects) for more than 6 months and obtained additional data. We collected their resting-state data and addiction features during these two times of scanning. All the data analyses steps were of the same to those in the cross-sectional analyses. We compared the post-pre test and observed a decrease of the recruitment in network community in those 22 IGD subjects (*t = -1*.*467, p = 0*.*095*), although the change did not reach the statistical significance. This suggests that continuous IGD situation further impaired their recruitment feature in executive control network. The result provides additional support to the conclusions.

## Discussion

Using the dynamic network analysis method, the current study provides a new perspective on the dynamic reconfiguration of the executive control network and reward network in subjects with IGD. The current study found a disturbed community structure (recruitment) in subjects with IGD, which may explain why subjects with IGD have impaired executive control abilities. Further analyses showed that the decreased recruitment coefficient was only observed in male subjects. This provides further support for previous findings on sex differences in IGD. We also explored the mediation effect between different networks.

### Subjects with IGD show disturbed community structure in the executive control network

‘Recruitment’ is the probability that a brain region is assigned to the same community as other nodes from its own network, and it is used to quantify the probability that a functionally-defined region of interest is assigned to the same community as functionally-defined regions of interest from the same subsystem ^17,18^. Generally, a reduced recruitment coefficient indicates that the nodes within the network are less likely to be divided into the same community over time.

In the current study, we learned from healthy controls that during the scanning period, the executive control process recruited stable regions within the executive control network and the reward network. Regions within these two systems exhibited a higher preference for intra-subsystem communication but a relatively low preference for inter-subsystem communication. However, in subjects with IGD, recruitment in the executive control network was unstable, and regions within the executive control network exhibited a lower preference for intra-subsystem communication but a relatively high preference for inter-subsystem communication. This suggests that subjects with IGD have an unstable executive network, and that the brain regions they recruit may differ each time they perform executive control. This disturbs the efficiency of executive control and prevents subjects with IGD from effectively controlling their impulses. For the reward network, there was no difference between the two groups.

Based on the patterns of recruitment features we observed, we concluded that subjects with IGD have a disturbed community structure in the executive control network; they had a lower preference for intra-subsystem communication but a relatively high preference for inter-subsystem communication. The decreased recruitment feature disrupted the efficiency of their executive control, which could explain why subjects with IGD have disrupted executive control and find it harder to control their gaming cravings.

### Disturbed community structure in male but not female subjects with IGD

Further analysis showed that the differences in recruitment between groups was only observed in male subjects with IGD, and no such features were observed in female subjects with IGD. As discussed in the introduction, studies have observed impaired executive control over reward seeking and craving for gaming ^1-3^; however, sex differences were observed in IGD studies. First, the prevalence of IGD is higher in males than in females, which suggests that males are more likely than females to develop IGD ^59^. At the same time, male subjects with IGD show more impaired executive control than female subjects with IGD, and it is harder for males to control their gaming cravings. The inhibitory control over game-elicited cravings in male subjects with IGD’ was more severely disrupted by gaming cues than that of females ^60^; short-term gaming elicited more craving-related activations to gaming cues in males vs. females ^61^. Brain regions implicated in executive control showed differential functional connectivity in males during gaming ^62^. Even for recreational gamers, female players exhibit better executive control than male players when facing gaming cues, which may provide resilience against developing IGD ^7^.

In the current study, decreased recruitment was only observed in male subjects with IGD, not females, and these findings provide further support to studies on sex differences in IGD. As we discussed on the role of recruitment, we can conclude that male subjects with IGD are less efficient in recruiting their executive control network regions, which provides an explanation for the underpinnings of the sex differences in executive control in IGD.

### The meditation effects suggest that subjects with IGD lack top-down control function

We also observed in RGUs that the integration coefficient of the RECN mediates the relationship between the recruitment of both the RECN and the BGN, which was not observed in subjects with IGD. The integration coefficient of the RECN reflected the dynamic interaction between nodes in the RECN and other networks. According to the framework of addictions, addiction is the result of an imbalance between the ECN and the BGN, which are generally considered to be structurally independent but functionally interacting ^63^. After an individual learns social rules, the ECN, which is primarily located in the frontal lobe, controls the subcortical BGN through several mechanisms, including decision-making and inhibitory control, which is considered to be the top-down control mechanism of the ECN in addiction ^64,65^. Under current conditions, we speculated that there was a stable top-down regulation mechanism based on a specific mediation framework in RGUs. However, this appears to be missing in subjects with IGD, which might be a potential risk factor for why IGD tends to be associated with dysfunctions of executive control.

### Limitations

Several limitations should be addressed. First, all subjects in the current study were free of comorbid disorders to exclude potential extra effects from other disorders. However, IGD is usually comorbid with other disorders, such as ADHD, nicotine addiction, or other types of psychiatric disorders. Second, all data were cross-sectional and lacked longitudinal data to verify the causal relationship between IGD and impaired executive control. Future studies should investigate this issue.

## Conclusions

First, subjects with IGD showed a lower recruitment coefficient than RGUs, making them less efficient at controlling their impulses. This could explain why subjects with IGD have impaired executive control over their gaming cravings. Second, decreased recruitment was observed only in male subjects with IGD, which is consistent with previous studies on sex differences in IGD. Third, the top-down regulation mechanism was only observed in RGUs but not in subjects with IGD, which suggests that subjects with IGD lack a stable top-down regulation mechanism. These conclusions revealed the mechanisms underlying why subjects with IGD exhibit impaired executive control over gaming cravings.

## Statistics and reproducibility

All statistical steps were used open software and we did not do any modification on them. The parameters were provided on each of the statistical steps.

## Acknowledgements

The current research was supported by The Cultivation Project of Province-levelled Preponderant Characteristic Discipline of Hangzhou Normal University (20JYXK008) and the Zhejiang Provincial Natural Science Foundation (LY20C090005).

This article has been posted on the preprint server BioRxiv.

## Author contributions

Min Wang designed this research and wrote the first draft of the manuscript. Min Wang and Hui Zheng analyzed the data and prepared the figures and tables and the longitudinal data analyses; Weiran Zhou, Lingxiao Wang, Shuaiyu Chen contributed to fMRI data collection, and manuscript revision. Guang-Heng Dong designed this research and edited the manuscript. All authors contributed to and approved the final manuscript.

## Data Availability

The data stored at our lab based network attachment system: http://QuickConnect.cn/others. ID:guests; PIN

