## supplementary materials for "Disrupted dynamic network reconfiguration of the executive and reward networks in internet gaming disorder: Why IGD subjects cannot control their gaming cravings"

### Statistical null models

In order to quantify the dynamic modular organization of the resting-state network in both IGD subjects and RGUs, we employed two random network null models: static null model and nodal null model (1, 2). The former created a multi-layer network with the same structure by randomly selecting the adjacency matrix and copying it 18 times. The latter creates a purely random connection model by randomly disrupting the correspondence between the nodes of each layer. Both test respectively the null hypothesis that network modular was static overtime, and the null hypothesis that the dynamic role of any node in the network is indistinguishable from the role of other nodes. For each subject, the null models were constructed by 100 times each, and 100 dynamic community detection iterations were performed on each generated empty model.

We respectively compared IGD and RGUs with their corresponding null models. The comparison index is $P_{ij}$ that corresponds to relative frequency that node $i$ and $j$ were assigned to the same community across time domain, also called module allegiance.

For the nodal null model, we compared the difference in mean module allegiance between the two groups and their null model. The real network module showed significantly higher regional interaction in the module allegiance in both group (IGD: $t = 3.247$, $p = 0.001$; RGU: t = 3.274, $p = 0.001$, see Fig.S1.A).

For the static null model, we compared the frequency distribution of module allegiance between the two groups and their null model. The real network module showed a distributed module allegiance values, while the distribution of the static null model is concentrated at 0 or 1 (see Fig.S1.BC; B: IGD group; C: RGU group).

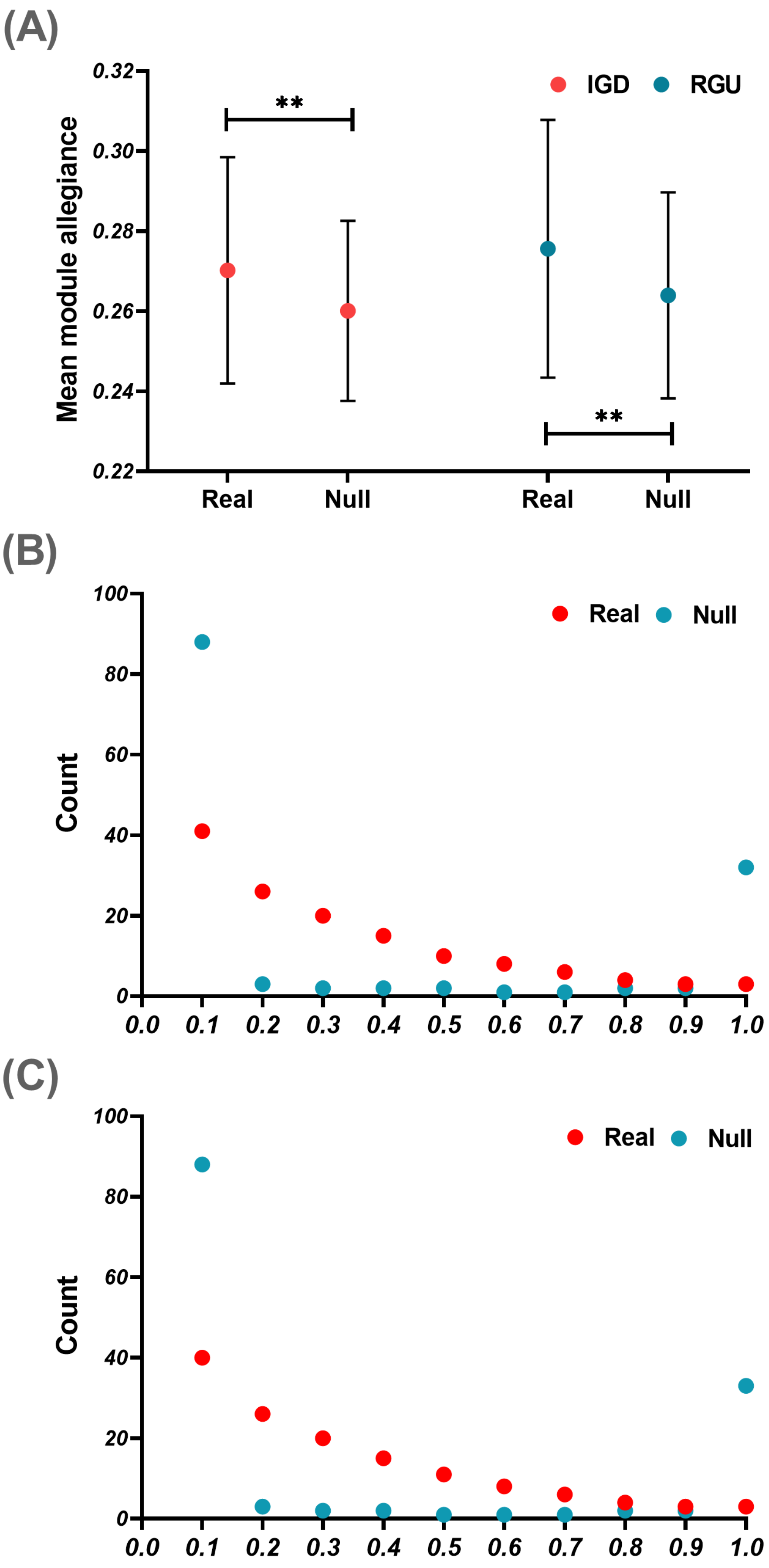
(A)
Mean module allegiance
0.32
0.30
0.28
0.26
0.24
0.22
IGD
RGU
**
**
Real
Null
Real
Null
(B)
Count
100
80
60
40
20
0
Real
Null
0.0
0.1
0.2
0.3
0.4
0.5
0.6
0.7
0.8
0.9
1.0
(C)
Count
100
80
60
40
20
0
Real
Null
0.0
0.1
0.2
0.3
0.4
0.5
0.6
0.7
0.8
0.9
1.0

**Fig.S1 Comparison of static and nodal null model**

Top: the comparison between the real network module and nodal null model in two group; Middle: the comparison between the real network module and static null model in IGD group; Bottom: the comparison between the real network module and static null model in RGU group. **: $p < 0.001$
